# Absence of subcerebral projection neurons delays disease onset and extends survival in a mouse model of ALS

**DOI:** 10.1101/849935

**Authors:** Thibaut Burg, Charlotte Bichara, Jelena Scekic-Zahirovic, Mathieu Fischer, Geoffrey Stuart-Lopez, François Lefebvre, Matilde Cordero-Erausquin, Caroline Rouaux

## Abstract

Amyotrophic Lateral Sclerosis (ALS) is a fatal neurodegenerative disease of adulthood that affects voluntary motricity and rapidly leads to full paralysis and death. ALS arises from the combined degeneration of motoneurons in the spinal cord and brain stem, responsible for muscle denervation, and corticospinal projection neurons (CSN), responsible for emergence of the upper motor neuron syndrome. Recent studies carried on ALS patients suggest that the disease may initiate in the motor cortex and spread to its projection targets. However, this “corticofugal hypothesis” of ALS has not yet been specifically challenged. Here, we provide a direct test of this hypothesis by genetically removing subcerebral projection neurons (SubCerPN), including CSN, in *Sod1^G86R^* mice, a mouse model of ALS. Ablation of the transcription factor *Fezf2*, leading to the complete absence of all SubCerPN, delays disease onset, reduces weight loss and motor impairment, and increases survival without modifying disease duration. Importantly absence of SubCerPN and CSN also limits pre-symptomatic hyperreflexia. Together, our results demonstrate that major corticofugal tracts are critical to ALS onset, and that SubCerPN and CSN in particular may carry detrimental signals to their downstream targets. In its whole, this study provides first experimental arguments in favour of the corticofugal hypothesis of ALS.

## Introduction

Amyotrophic lateral sclerosis (ALS) is a devastating neurodegenerative disease characterized by rapidly progressing muscle atrophy and paralysis, leading to death within only two to five years of diagnosis. Clinically and histologically, ALS is defined as the combined and progressive loss of two broad neuronal populations involved in motor control: the corticospinal and corticobulbar neurons (CSN, or upper motor neurons) in the motor cortex, and the spinal and bulbar motoneurons (MN, or lower motor neurons) in the brain stem and spinal cord [8,63]. This duality of targeted neuronal populations and affected regions of the central nervous system has fostered a long-time debate regarding the origin of the disease along the cortico-spino-muscular axis (reviewed in [54]).

The French Neurologist Jean-Martin Charcot who provided the first description of ALS suggested a cortical origin of the disease and a descending propagation from the motor cortex to the spinal cord [11]. Following Charcot’s histological description of the disease, comprehensive clinical examination of ALS patients unravelled series of signs highly suggestive of a cortical origin of the disease, such as the split hand syndrome and typical gait abnormalities for instance [19]. Meanwhile, trans-cranial magnetic stimulation studies unravelled early hyperexcitability of the motor cortex that characterizes both sporadic and familial ALS patients, negatively correlates with survival, and manifests prior to disease onset, pointing to a potential role of the motor cortex in disease initiation [67]. Cortical hyperexcitability has been proposed to translate into glutamatergic excitotoxicity to the downstream targets of the corticospinal neurons, the alpha motoneurons of the brain stem and spinal cord, providing a first possible mechanism for corticospinal propagation [18]. More recently, the proposed staging of TDP-43 pathology [6], a histopathological burden that characterizes the vast majority of ALS patients, led to the emergence of the so-called corticofugal hypothesis. According to it, the disease may originate in the motor cortex and misfolded TDP-43 proteins may disseminate to the direct, mono-synaptic targets of cortical projection neurons along the main corticofugal routes with a prion-like mechanism [5]. If corticofugal propagation is supported by longitudinal diffusion tensor imaging studies with tractography or connectome analysis of ALS patients [32,66], it cannot be directly tested in patients. In spite of the heavy differences that exist between species, and in particular the uniqueness of the Human and primate corticomotoneuronal system [18], we reasoned that mouse genetics could prove useful to directly address a potential cortical origin and corticofugal propagation of ALS.

In rodents, like in Humans, corticofugal projections arise from two broad populations of excitatory projection neurons, *i.e.*, the subcerebral projection neurons (SubCerPN) and the corticothalamic projection neurons (CThN) [41]. CThN are located in the cortical layer VI and connect the cerebral cortex to numerous thalamic nuclei, that connect back to the cerebral cortex, creating a loop of reciprocal sensory processing [7]. SubCerPN are located in the cortical layer V and connect the various areas of the cerebral cortex to their most distant targets. While the thalamus is affected by the TDP-43 pathology as early as stage 2 [6], and shows impairment in imaging studies [14], no major contribution of CThN to ALS has been suspected. SubCerPN instead include, amongst other subpopulations, the disease-relevant corticospinal neurons (CSN) that connect the cerebral cortex to the spinal cord. While major differences exist between primates and rodents regarding the route of the corticospinal tract within the spinal cord, and the connectivity mode of the CSN onto the alpha motoneurons, *i.e.* both mono- and pluri-synaptic in primates, and essentially pluri-synaptic in adult mice [36], it is worth highlighting that many mouse models of the disease recapitulate CSN or SubCerPN degeneration [22,28,38,42,50,70,71]. In this regard, we recently showed that *Sod1^G86R^* mice not only display pre-symptomatic CSN degeneration, but also that CSN and spinal motoneuron degeneration are somatotopically related [42], similarly to what had been described in ALS patients [46,53,72].

Here, we sought to test the contribution of CSN and other SubCerPN to disease onset and progression by taking advantage of the *Fezf2* knock-out mice [29] that develop in absence of SubCerPN. These animals lack the gene encoding a master transcription factor that is both necessary [12,13,39,40,48] and sufficient to instruct the birth and specification of CSN and other SubCerPN from embryonic cortical progenitors [12,39,48] but also from non-cortical neural progenitors and even post-mitotic neurons of another lineage [57,58]. We thus crossbred the *Sod1^G86R^* and the *Fezf2^−/−^* mouse lines to generate a model that ubiquitously expresses a mutant of the murine *Sod1* gene, a condition sufficient to develop ALS-like symptoms and premature death [16,42,56,59], but entirely lacks CSN and other SubCerPN, hence challenging ALS definition.

## Materials and Methods

### Animals

All animal experiments were performed under the supervision of authorized investigators and approved by the local ethical committee of Strasbourg University (CREMEAS, agreements # 00738.01). Animals were housed in the animal facility of the Faculty of Medicine of Strasbourg, with a regular 12 hours light/dark cycle, under constant conditions (21±1°C; 60% humidity). Standard soft laboratory food and water were accessible ad libitum throughout all experiments. BAC transgenic *Sod1^G86R^* mice [56] were obtain from the animal facility of the Faculty of Medicine of Strasbourg, and knockout *Fezf2^−/−^* mice were generated by Hirata and colleagues [29] and kindly provided by the Arlotta Lab. Mice were genotyped by PCR of genomic DNA from tail biopsies as previously described [29,56]. *Sod1^G86R^* males were crossed with *Fezf2^+/−^* females and F1 *Fezf2^+/−^/Sod1^G86R^* males were crossed *Fezf2^+/−^/WT* females. The F2 generation provided four genotypes of interest: *Fezf2^+/+^/WT; Fezf2^+/+^/Sod1^G86R^; Fezf2^−/−^/WT; Fezf2^−/−^/Sod1^G86R^*. Males were used for survival and behavioural studies and end-stage histology. Males and females, in equal proportions, were used for H-reflex and histology at pre-symptomatic ages. Mice were followed daily, and disease progression was rated according to a clinical scale going from score 4 to 0, as previously described [42]. End-stage animals were euthanized upon reaching score 0, *i.e.* when they were no longer unable to roll over within 10 s after being gently placed on their back [42]. Disease onset was calculated as the time of peak of body weight.

### qPCR analyses

Cortex and spinal cord were harvested and rapidly frozen in liquid nitrogen and stored at - 80°C until analysis. Total RNA was extracted using TRIzol reagent (Invitrogen) and stainless-steel bead in a Tissue Lyser (Qiagen). 1 µg of RNA was reverse transcribed using the iScript cDNA synthesis kit (Bio-Rad). Quantitative PCR (qPCR) was performed with the IQ SYBR Green Supermix (Bio-Rad). Gene expression was normalized by calculating a normalization factor using *Gusb*, *Actb* and *Hsp90ab1* as reference genes. The following primer sequences were used for qPCR:

> *Gusb*: F-CGAGTATGGAGCAGACGCAA; R-AGCCTTCTGGTACTCCTCACT
>
> *Actb*: F-ATGTGGATCAGCAAGCAGGA; R-AGCTCAGTAACAGTCCGCCT
>
> *Hsp90ab1*: F-TACTACTCGGCTTTCCCGTCA; R-CCTGAAAGGCAAAGGTCTCCA
>
> *Fezf2*: F-GTGCGGCAAGGTGTTCAATG; R-CAGACTTTGCACACAAACGGT
>
> *Sod1*: F-GAGACCTGGGCAATGTGACT; R-GTTTACTGCGCAATCCCAAT
>
> *ChAT*: F-CTGGCCACCTACCTTCAG-TG; R-CCCCAAACCGCTTCACAATG

### Motor tests and regression analyses

For all motor tests, mice were trained from 5 to 8 weeks of age, and then followed from 9 weeks of age until death, on a daily basis for general health and neurological symptoms, twice a week for body weight and muscle grip strength, and once a week for inverted grid test, accelerating rotarod and CatWalk. Each motor session consisted of three trials and the results represent the mean of these three trials. Motor coordination and endurance were assessed using rotarod (Ugobasile model 7650), with trials of 300 seconds composed of an acceleration period of 150 seconds (4-20 rpm) followed by constant speed period of 150 seconds. Muscle strength was measured using a strength grip meter (Bioseb, BIO-GS3). Four limbs hang test that allows measuring the ability of mice to use sustained limb tension to oppose gravitational force was used as previously described [10,60]. Briefly, mice were placed on a cage grid, allowed to accommodate to their environment for 3-5 seconds before the grid was slowly inverted and positioned 35 cm above a receiving cage filled with 5-6 cm of bedding. The hanging time corresponds to the time the mice spent hung to the grid before dropping it. Mouse gait was analysed with the CatWalk XT (Noldus Information Technology). This automated gait analysis system allows digitalizing each individual footprint during the mouse locomotion and generates numerous parameters for quantitative and qualitative analysis of individual footprint and gait [65] (Supplementary Table 1). Recordings were performed under the conditions previously described [60]. Each mouse was allowed to cross freely the recording field of the runway with three independent attempts. Criteria for data collection were *i)* crossing the field in less than 10 seconds and *ii)* a walking speed variation of less than 60%. Multiple linear regression analyses were run on R 3.4.3 with all the relevant packages. They allowed including weight only (Grip Test, Rotarod, Inverted Grid) or weight and speed (all Catwalk parameters except for the speed itself) as confounding variables. Normality of the distributions was tested using the Shapiro-Wilk test or the Kolmogorov-Smirnov test, and was assessed graphically using a normal quantile plot.

### Electromyography

All recordings were performed with a standard EMG apparatus (Dantec) on mice anesthetized with a solution of Ketamine (Imalgène 1000®, Merial; 90mg/kg body weight) and Xylazine (Rompun 2%®, Bayer; 16mg/kg body weight) and kept under a heating mat to maintain physiological muscle temperature (≈31°C). Electrical activity was monitored on muscles of both limbs for at least 2 minutes, as previously described [59].

### Tail spasticity

Tail spasticity of end stage mice was determined and quantified as previously described [20]. Briefly, a monopolar needle electrode (Medtronic, 9013R0312, diameter 0.3 mm) was inserted in segmental tail muscles of paralyzed mice to record reflex activity. Muscles spasms were evoked with mechanical stimulation of the tail. For quantification, signal intensities were measured before and after stimulation, using ImageJ (NIH), and the signal-to-noise ratio was calculated.

### H-reflex recording

H-reflex was assessed on both hindlimbs (with a three weeks interval) of pre-symptomatic mice aged of 80 and 105 days, by modifying a previously described method [35]. Mice were anesthetized with a solution of Ketamine (Imalgène 1000®, Merial; 90mg/kg body weight) and Xylazine (Rompun 2%®, Bayer; 16mg/kg body weight) and placed on a heating pad. After unilateral exposure of the sciatic nerve, a spherical homemade stimulation electrode was placed around the nerve. A recording monopolar needle electrode (Medtronic, 9013R0312, diameter 0.3 mm) was transcutaneously placed in the abductor digiti minimi muscles and recordings were obtained with an amplifier (Digitimer Ldt, DS3). A first stimulation of 0.2 ms at 0.1 Hz was initially applied, followed by gradually increasing current intensity. This protocol allowed us to find the minimal intensity to elicit a M-response and the intensity to obtain a maximal M-response, in order to fix the stimulation intensity at approximately 50% of the M-response. For each animal, we tested the presence or absence of the H-reflex by performing 10 sciatic nerve stimulations at 0.1Hz, 0.2Hz, 0.5Hz and 1Hz. For each of the 10 traces, we measured the maximal amplitude (peak value – through value) of the noise, on a 1 ms window between stimulation and M-Wave (ΔNoise). We measured similarly the amplitude of the H-Reflex as the peak-through value of the recording on a 1 ms window situated at the expected latency for the H-reflex, between 5 and 8 ms following stimulation (ΔResponse). The mean and standard deviation of the ΔNoise were measured from the 10 ΔNoise values obtained from the 10 successive traces for each animal, and used to calculate the Z-scores. The Z-score of each response to the stimulation site was calculated using the following equation:

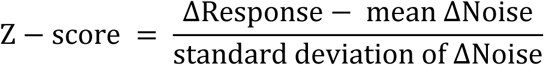

A Z-score of 3, corresponding to a significance level of 0.001, was chosen to discriminate significant responses from non-significant ones. This conservative threshold was chosen in agreement with visual inspection of the traces, to ensure that any H-reflex response was not simply due to noise. H-reflex and F-waves share some electrophysiological properties, but F-wave’s persistence, defined as the number of significant responses divided by the number of stimuli, is mainly independent of stimulation frequency, or only tends to increase above 1-2Hz stimulations rates [21,43]. To ensure that the considered signal was an H-reflex, we thus measured the persistence of the response across increasing stimulations rates, and only considered signals not demonstrating persistence (Supplementary Fig.2). After recording, mice were stitched and kept under a heating lamp until recovery. Mice were then monitored for recovery and subjected to daily checks. Mice were considered to display H-reflex when at least one of the two legs was positive.

### Retrograde labelling of the CSN

CSN were labelled as previously described [42]. Briefly, animals were deeply anesthetized with an intraperitoneal injection of Ketamine (Imalgène 1000®, Merial; 120 mg/kg body weight) and Xylazine (Rompun 2%®, Bayer; 16 mg/kg body weight) solution and placed on a heating pad. Laminectomy was performed in the C3-C4 cervical region of the spinal cord, and the dura was punctured using a pulled glass capillary lowered to the dorsal funiculus. Five pressure microinjections of 23 nl of Fluorogold (Fluorchrome) were performed on each side of the dorsal funiculus (Drummond Scientific, Nanoject II). Five days after injection, mice followed the histological procedure (see below).

### Histological procedures

Animals were deeply anesthetized with an intraperitoneal injection of Ketamine (Imalgène 1000®, Merial; 120 mg/kg body weight) and Xylazine (Rompun 2%®, Bayer; 16 mg/kg body weight), and transcardially perfused with cold 0.01 M PBS, followed by cold 4% PFA in 0.01 M PBS. Brains, spinal cords, gastrocnemius and tibialis were post-fixed in the same fixative solution overnight (nervous tissues) or for 1 hour (muscles) and stored in PBS 0.1M at 4°C until use. Fixed brains and spinal cords were cut in 40µm thick sections on a vibratome (Leica Biosystems, S2000). Fluorogold-labelled brains were mounted on slides with DPX mounting solution (Sigma-Aldrich). Immunostainings on brains and spinal cords were performed on slides at room temperature. First, sections were immersed in 3% hydrogen peroxide (H_2_O_2_) to remove the endogenous peroxidase activity and washed with phosphate buffered saline (PBS). Unspecific binding sites were blocked with 5% horse serum and 0.5% triton X-100 in PBS for 30 min and incubated with primary antibody overnight. After rinsing in PBS, sections were incubated biotinylated secondary antibody for 2 hours, rinsed in PBS and incubated 1 hour with the Vectasatin ABC Kit (Vector Laboratories, PK7200). Revelation was performed by incubating the sections in 0.075% 3, 3’-diaminobenzidine tetrahydrochloride (Sigma-Aldrich, D5905) and 0.002% H_2_O_2_ in 50mM Tris HCl. Lumbar (L2-L3) spinal motor neurons from six sections covering a distance of 640 µm were immunolabelled using a goat anti-choline acetyltransferase (ChAT) antibody (Merk Millipore, AB144P, 1/50) and a biotinylated donkey anti-goat IgG (Jackson, 1/500). Antibody for other spinal cord immunostainings were: rat anti CTIP2 (Abcam, 18465, 1/100), mouse anti P62 (Abcam, 56416, 1/100), rabbit anti GFAP (Dako, Z0334, 1/200), goat anti Iba1 (Abcam AB5076, 1/100) and corresponding biotinylated secondary antibody (Jackson, 1/500). Serotonergic neurons of the median and dorsal raphe nuclei from individual coronal sections located at Bregma - 4.72 mm were immunolabelled with a goat anti-TPH2 antibody (Abcam, AB121013, 1/500), followed by a donkey anti-goat biotinylated secondary antibody (Jackson, 1/500). Tibialis anterior muscles were dissected into bundles and processed for immunofluorescence using a combination of rabbit anti-synaptophysin and rabbit anti-neurofilament antibodies (Eurogentec) followed by an Alexa-conjugated donkey anti-rabbit 488 (Jackson, 1/1 000), and rhodamine conjugated α-bungarotoxin (Sigma-Aldrich, T0195), as previously described [42]. On average 100 NMJ per animals were examined. Images were captured using an AxioImager.M2 microscope (Zeiss) equipped with a structured illumination system (Apotome, Zeiss) and a high-resolution B/W camera (Hamamatsu), and run by the ZEN 2 software (Zeiss). Experimenters blinded to the genotypes performed the neuronal counts and NMJ assessments.

### Statistical Analysis

Data are presented as mean ± standard error of the mean (SEM). Statistical analysis was performed using GraphPad Prism 6 (GraphPad, CA). For comparison between two groups, Student’s t-test was used and for three or more groups, 1-way analysis of the variance followed by Tukey’s multiple comparison post hoc test was applied. If there were 2 variables, 2-way analysis of the variance followed by Tukey’s multiple comparison post hoc test was applied. For survival and disease onset analysis, animals were evaluated using log-rank test (Mantel-Cox). Grubbs’test or ESD method (extreme studentized deviate) has excluded one mouse of our survival experiment. Fischer exact test was used to determine significant different proportion. Results were considered significant when p < 0.05.

## Results

### Generation of the *Fezf2^−/−^*; *Sod1^G86R^* mice

To test the contribution of the motor cortex and more broadly the cerebral cortex to ALS onset and progression, we generated a mouse line that overexpresses the mutant murine *Sod1^G86R^* transgene but lacks one of the most important cortical output, the SubCerPN. We crossbred *Sod1^G86R^* mice, a well-established model of ALS [16,56,59], to mice knocked out for the gene encoding the transcription factor *Fezf2* and that develop in absence of cortical layer V SubCerPN [29,48] (Fig. 1a). As previously reported, retrograde labelling of CSN, a subpopulation of SubCerPN, by injection of Fluorogold in the cervical portion of the dorsal funiculus, in the spinal cord of 75 day-old animals, revealed labelled pyramidal cells in the layer V of the motor area of *Fezf2^+/+^* (*WT*) mice but not in that of *Fezf2^−/−^* (*KO*) mice (Fig. 1b). We ran two steps of crossbreeding to obtain four genotypes of interest: *Fezf2^+/+^* and non-transgenic (*WT*), *Fezf2^−/−^* and non-transgenic (*KO*), *Fezf2^+/+^* and *Sod1^G86R^* (*Sod1*), and *Fezf2^−/−^* and *Sod1^G86R^* (*KO/Sod1*) (Fig. 1a). To further verify the absence of SubCerPN in *KO* and *KO/Sod1* animals, we performed a fluorescent immunostaining to reveal CTIP2, a transcription factor expressed in the large nuclei of layer V SubCerPN, and in the smaller nuclei of layer VI CThN [1]. Microscopic analysis of coronal sections revealed the absence of CTIP2-positive layer V SubCerPN in *KO* and *KO/Sod1* animals compared to their *WT* and *Sod1* littermates (Fig. 1c). To check that absence of *Fezf2* expression and of SubCerPN had no effect on *Sod1* expression or on spinal motoneurons (MN) generation and specification, we ran qPCR analysis in the cerebral cortex and lumbar spinal cord of 75 day-old animals (Fig. 1d). Up-regulation of *Sod1* expression was verified in *Sod1* and *KO/Sod1* animals and did not differ between the two genotypes, neither in the cerebral cortex nor in the spinal cord (Fig. 1d). Finally, expression of the motoneuronal marker *Chat* was not significantly different between any of the four genotypes, ruling out the possibility that absence of SubCerPN, and more particularly of CSN may have a detrimental effect on spinal MN birth and specification (Fig. 1d), in accordance with the fact that *Fezf2* is not expressed in the spinal cord [29,48]. Together, the data suggest that *KO/Sod1* mice may represent a good model to study the impact of SubCerPN on ALS-like onset and progression, and to test the corticofugal hypothesis.

**Fig. 1.**
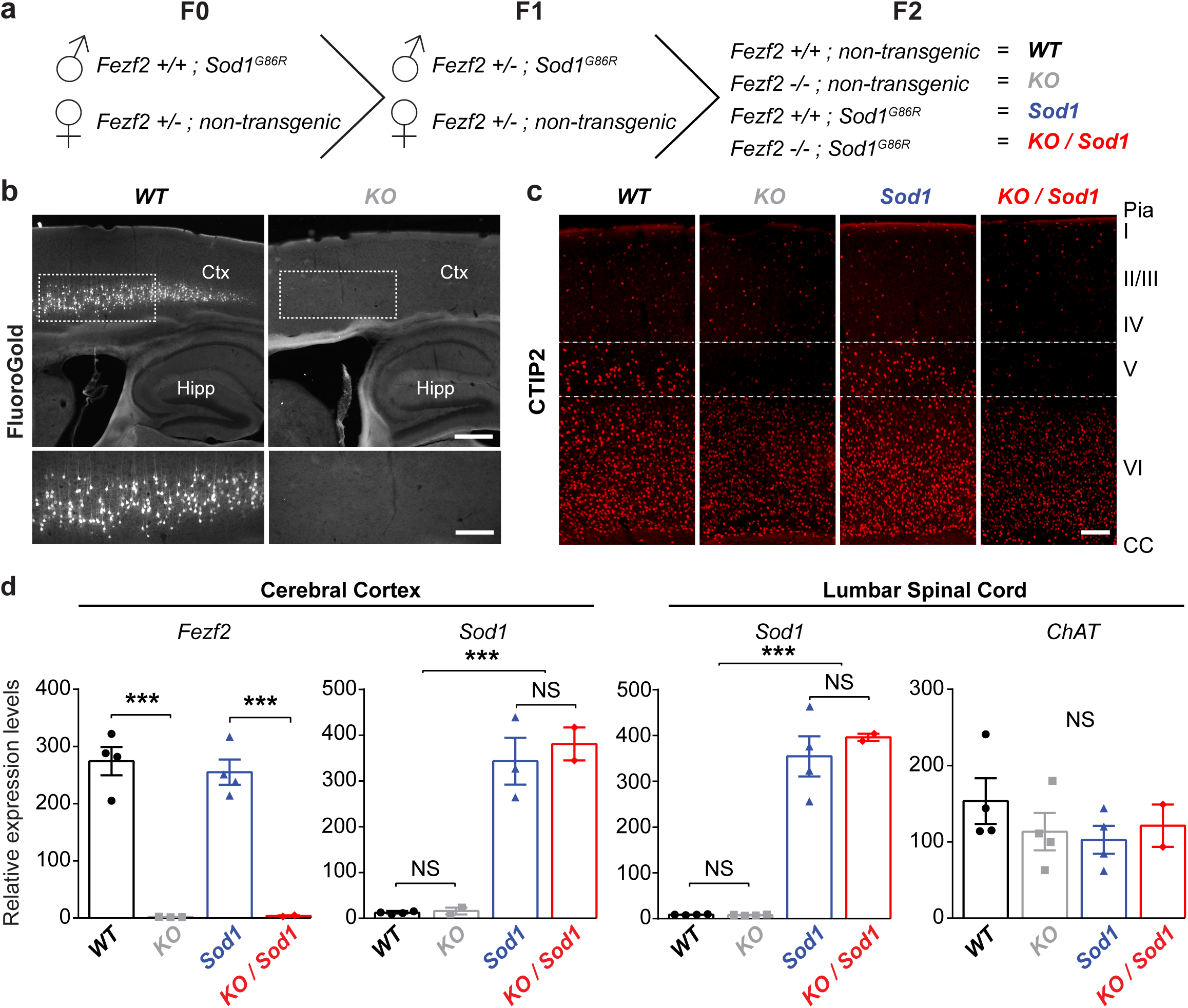
Generation of a mouse model overexpressing the *Sod1^G86R^* transgene but lacking all SubCerPN, including the CSN. **a** Schematic of the crossbreeding of the *Sod1^G86R^* and *Fezf2^−/−^* mouse lines to generate four genotypes of interest: *WT, KO, Sod1 and KO/Sod1*. **b** Retrograde labelling of the CSN from the spinal cord of *KO* animals (right) and *WT* (left) showing absence of CSN in *KO* mice. N = 5 for all genotypes. **c** Representative images of brain coronal sections, at the level of the motor cortex, showing CTIP2 immunolabelling of layer V SubCerPN and of layer VI CThPN in *WT* and *Sod1* mice, and confirming the absence of cortical layer V SubCerPN from the *KO* and *KO/Sod1* animals. N = 3 for all genotypes. **d** qPCR analysis of *Fezf2*, *Sod1* and *Chat* expression in the cerebral cortex and spinal cord indicating that absence of *Fezf2* does not affect *Sod1* or *Chat* expression; 2-way ANOVA; N = 4 *WT*, 4 *KO*, 4 *Sod1*, 2 *KO/Sod1*; *** p < 0.001; Scale bar = 400 µm in upper panels and 200 µm in lower panels of **b**, and 250 µm in **c**.

### Absence of SubCerPN delays disease onset and death

To examine the consequences of the absence of projections from the cerebral cortex to its main subcerebral targets, we assessed and compared general clinical parameters of *Sod1* and *KO/Sod1* mice and their controls. Weight loss is a hallmark of ALS and negatively correlates with survival in ALS patients [17]. Similarly, *Sod1^G86R^* mice start losing weight before onset of motor deterioration, and weight maintenance is positively correlated with their survival [16]. *Fezf2* deficient mice present a smaller weight gain compared to wild type mice [29], a condition that could *per se* be detrimental in a context of ALS [17]. Weight analysis of 75 day-old animals confirmed that *KO* mice were indeed lighter than *WT* mice, and revealed, as expected, that *KO/Sod1* were also lighter than *Sod1* animals. Body mass index analysis on another cohort of animals revealed that reduced weight was associated with reduced size and that *KO* and *KO/Sod1* mice were smaller but not thinner than their *WT* and *WT/Sod1* littermates (data not shown). At this age, we observed no difference either between *Sod1* and *WT* or between *KO/Sod1* and *KO* mice (Fig. 2a). Long-term weight follow-up showed that both *WT* and *KO* mice regularly gained weight, but that *Sod1* and *KO/Sod1* mice instead stopped gaining weight prematurely (Fig. 2b). This arrest of weight gain, or peak of weight, is often used a measure of disease onset in ALS mouse models [4,16]. *KO/Sod1* mice presented a significant delay of disease onset compared to their *Sod1* littermates (Median: 107.5 days vs 159 days; p = 0.0070) (Fig. 2c). Similarly, the survival of the *KO/Sod1* mice was increased compared to that of *Sod1* mice (Median: 130 days vs 183 days; p = 0.0083) (Fig. 2d), but the overall disease duration was not significantly different between the two genotypes (Fig.2e). Finally, the weight loss during the course of the disease was significantly smaller for the *KO/Sod1* mice than it was for the *Sod1* mice (36.45 ± 1.82 % vs 25.07 ± 1.68 %; p < 0.0001) (Fig.2f), in accordance with an arrest of weight gain rather that a clear weight loss, as seen also on the longitudinal weight follow-up (Fig. 2b), and an increased survival. Together, the data indicate that, in the *Sod1^G86R^* mouse model of ALS, absence of SubCerPN is beneficial as it delays disease onset and death, and limits weight loss during the course of the disease.

**Fig. 2.**
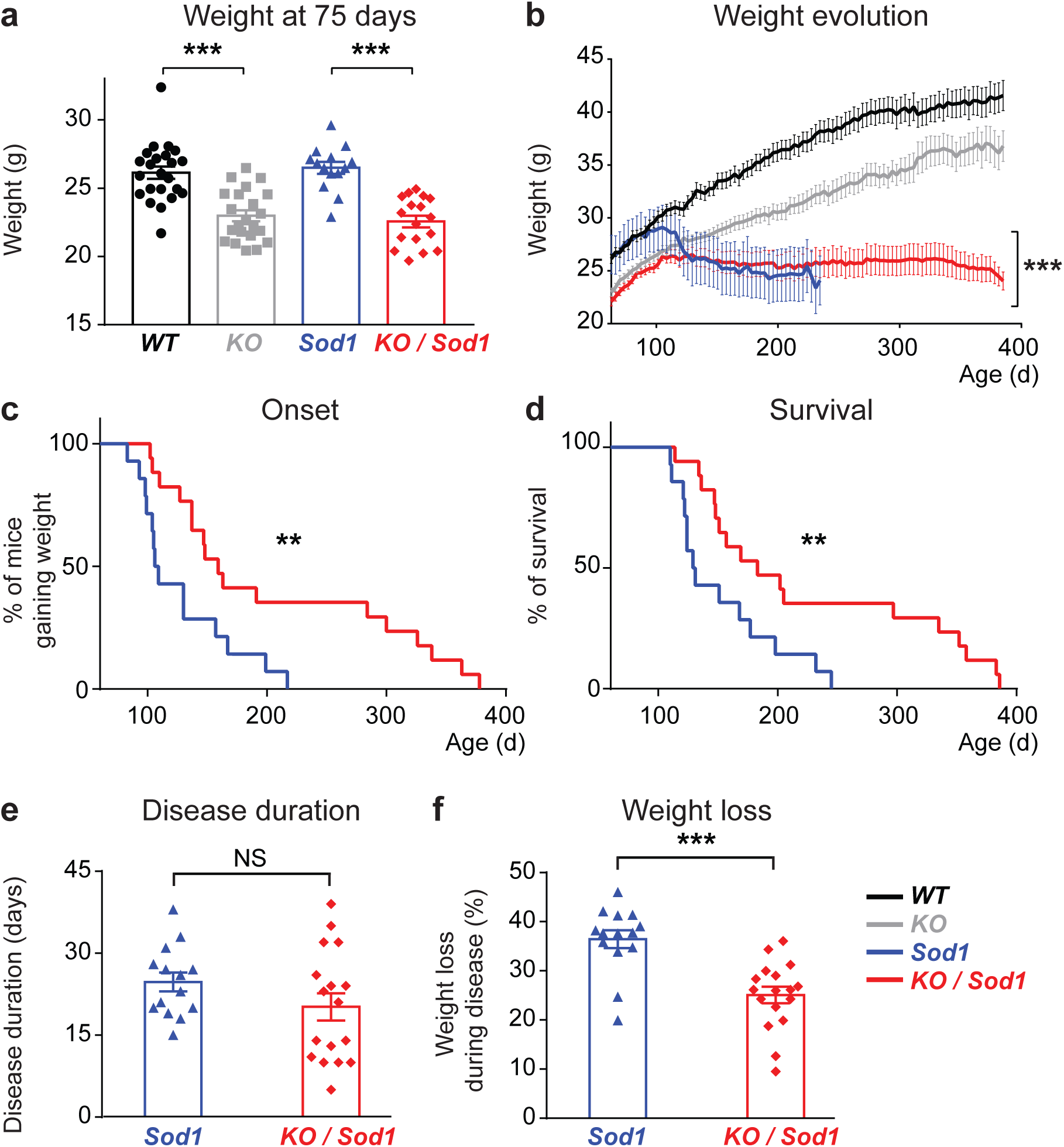
Absence of SubCerPN delays onset, prevents weight loss, and prolongs survival of *Sod1^G86R^* mice. **a** Bar graph representing the average weight of the four groups of mice at the beginning of the survival study (75 days); 1-way ANOVA. **b** Graphical representation of the evolution of the weight over time for the four genotypes. Note that while *Sod1* mice clearly lost weight, *KO/Sod1* mice rather stopped gaining weight; linear mixed effects model. **c-d** Kaplan-Meier plots of disease onset, define as the time when animals stopped gaining weight (**c**), and survival (**d**) in days, for *Sod1* and *KO/Sod1* mice; log-rank test (Mantel-Cox). **e-f** Bar graphs representing disease duration in days (**e**) and the percentage of weight loss during the course of the disease (**f**); Student unpaired t-test. For all data, N = 23 *WT*, 22 *KO*, 14 *Sod1*, 17 *KO/Sod1*; ** p < 0.01; *** p < 0.001.

### Absence of SubCerPN attenuates motor impairment

We next tested whether absence of SubCerPN could also ameliorate motor performances, and more particularly muscle strength, assessed by grip and inverted grid tests, motor coordination, assessed on accelerating Rotarod, and gait, assessed by CatWalk (Fig. 3 and 4). Because *KO* and *KO/Sod1* mice displayed initially a smaller weight, and because *Sod1* and *KO/Sod1* lost weight during the course of the disease, we ran linear regression analysis using the weight as a co-variable in order to better compare the performances of the different groups of animals (Fig. 3g-i and Supplementary Table 1). We ran two series of three comparisons: *WT* vs *Sod1*, *WT* vs *KO*, and *KO* vs *KO/Sod1*, both at the time when the experiment was initiated (Origin, Supplementary Table 1) and over the course of the experiment (Fig. 3g-i, and Slope, Supplementary Table 1). On the grip test and Rotarod, *WT* and *KO* mice displayed similar performances and maintained them over time (Fig. 3a,b). *Sod1* and *KO/Sod1* mice instead rapidly showed decreased performances (Fig. 3a,b and g,h), but *KO/Sod1* mice were affected later than their *Sod1* littermates (Fig. 3d,e) and maintained higher performances over time than their *Sod1* littermates (Fig. 3a,b and 3g,h). On inverted grid, *KO* and *KO/Sod1* mice initially displayed more difficulty in opposing their gravitational force compared to *WT* and *Sod1* mice, and this in spite of their smaller weight (Fig. 3c and i). Yet, while *WT* and *KO* animals maintained their hanging time throughout all the repetitive assessments, *Sod1* and *KO/Sod1* saw their performances gradually decreasing (Fig. 3c,i). This decrease occurred later in *KO/Sod1* animals than in *Sod1* animals (Fig. 3f). In addition, the impairment rate of *KO/Sod1* was milder than that of *Sod1* (Fig. 3c,i), and, by disease end stage, the *KO/Sod1* performed better than the *Sod1* (Fig. 3c,i). Linear regression analyses of the grip strength, and inverted grid test, indicated that significant differences existed between the *WT* and *KO* animals initially (Origin, Supplementary Table 1, p < 0.001), but that over time they evolved in a similar manner (Fig. 3g,i and Slope, Supplementary Table 1, NS). Such differences were expected given that absence of SubCerPN implies absence of corticospinal neurons and discrete modifications of the motor behaviour in mice, without evolution over time. Importantly, linear regression analyses of the grip strength, the Rotarod and the inverted grid data, over time, indicated significant differences between *KO* and *KO/Sod1* animals further confirming that absence of SubCerPN moderated motor deterioration in mice (Fig. 3g-i).

**Fig. 3.**
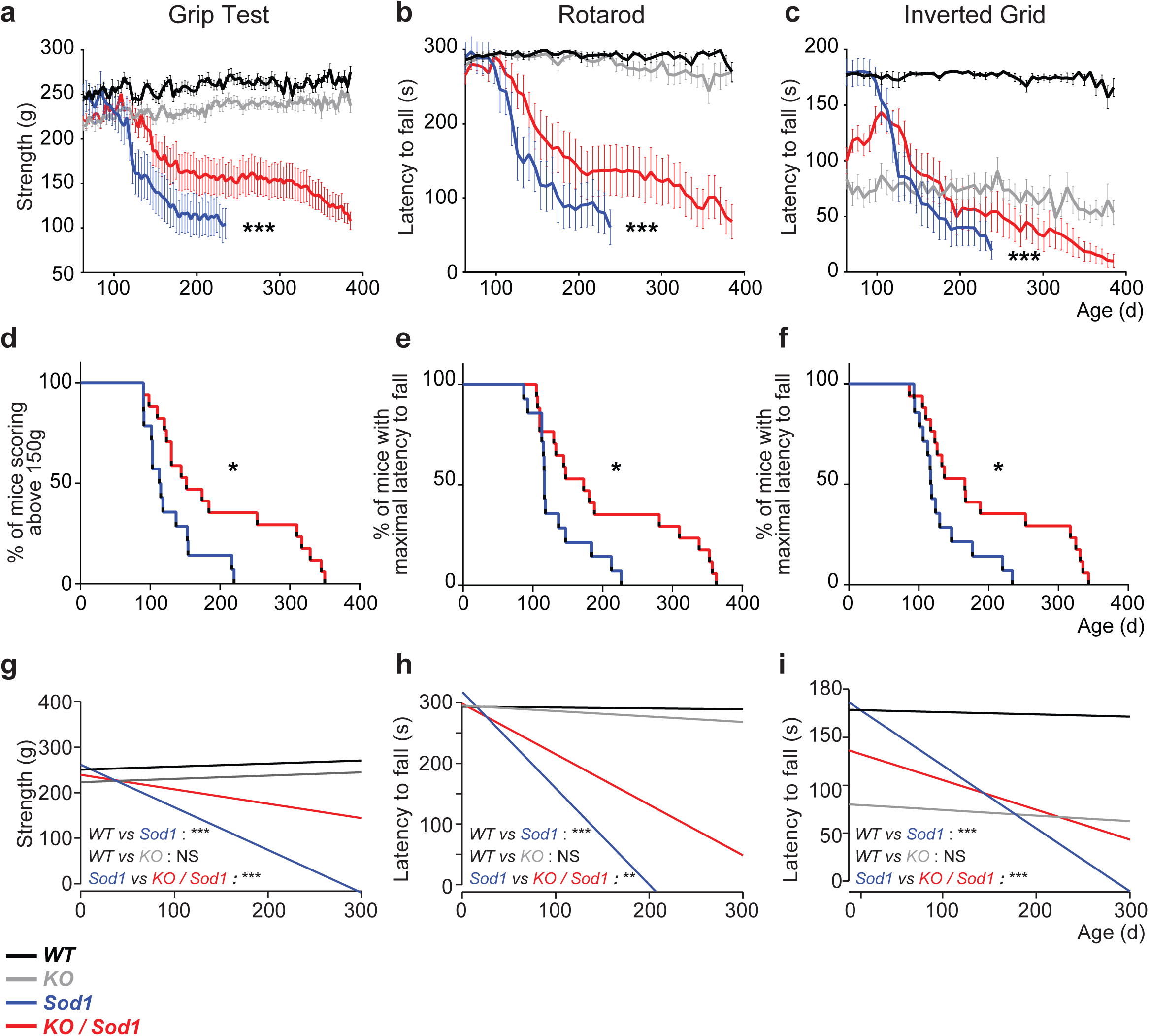
Absence of SubCerPN delays onset of motor symptoms and slows the decline of motor capacities. **a-c** Graphical representation of motor capacities over time, in days, on the grip strength test (**a**), the rotarod test (**b**) and the inverted grid test (**c**); linear mixed effects model. **d-f** Kaplan-Meier plots of grip strength score above 150g (**d**), maximal latency to fall from the rotarod (**e**) or from the inverted grid (**f**); log-rank test (Mantel-Cox). **g-i** Linear regression analysis conducted using the weight as covariate; the comparisons represented here are those of the slopes. For all data, N = 23 *WT*, 22 *KO*, 14 *Sod1*, 17 *KO/Sod1*; * p < 0.05, ** p < 0.01, *** p < 0.001, NS: non-significant.

**Fig. 4.**
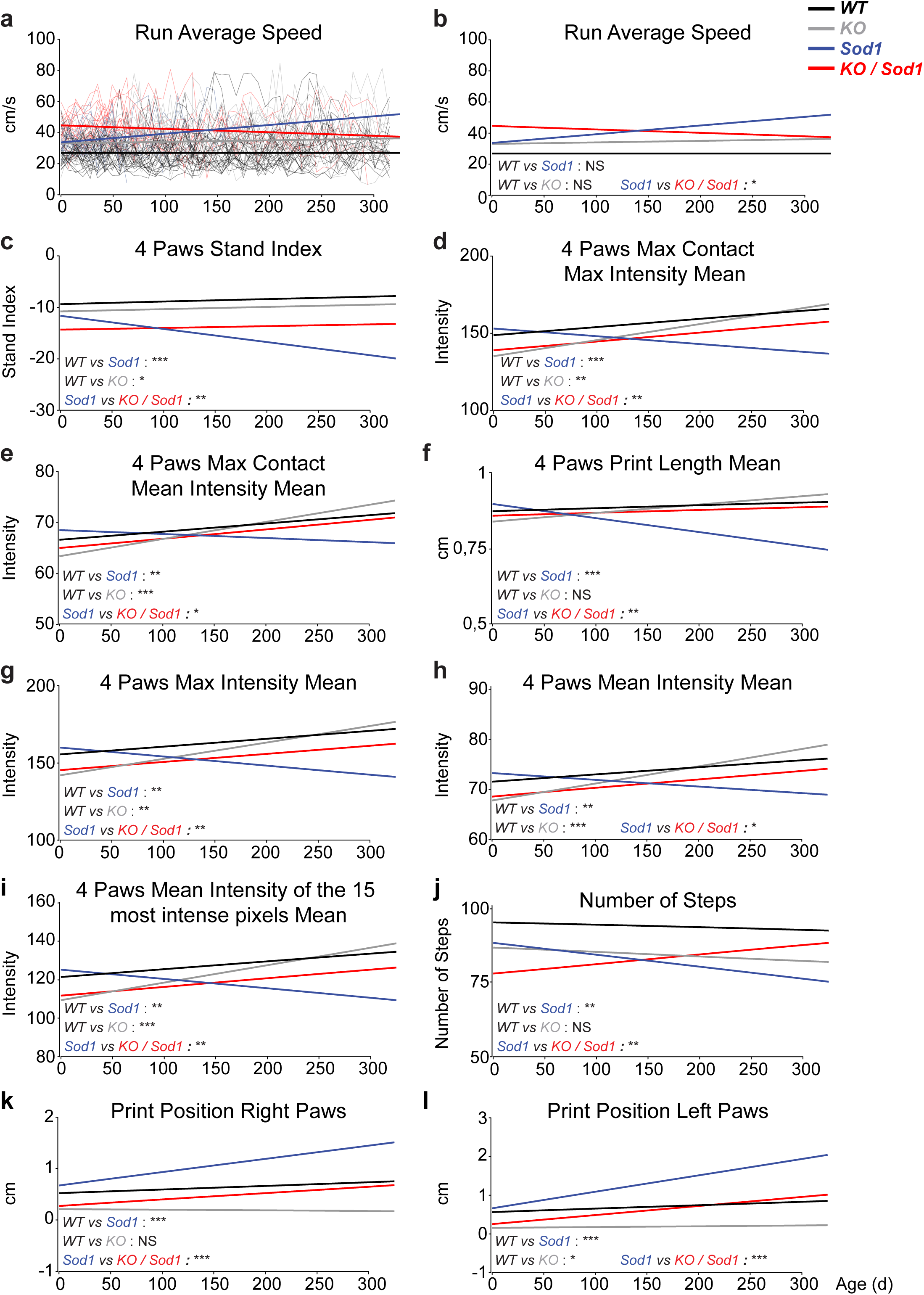
Absence of SubCerPN ameliorates gait parameters recorded on CatWalk. **a-l** Linear regression analyses were used to model overall evolution of the groups of animals taking into account the individual progressions of each mouse over time. Weight was used as a covariate in **a-b** and weight and speed were used as covariates in **c-l**. Individual mouse traces were removed (**b-l**) to ease visualization of the regression curves (compare **b** to **a**). The comparisons represented here are those of the slopes. For all data, N = 23 *WT*, 22 *KO*, 14 *Sod1*, 17 *KO/Sod1*; * p < 0.05, ** p < 0.01, *** p < 0.001, NS: non-significant.

CatWalk gait analysis device allows testing more than 200 parameters that can be classified into five broad categories: run characterization, temporal, spatial, kinetic and inter-limb coordination parameters [9]. We ran linear regression analyses on the CatWalk data, using not only the weight as a covariate, but also the speed, already shown to greatly influence numerous gait parameters [2] (Fig. 4 and Supplementary Table 1). We identified 3 spatial parameters that were significantly different between *WT* and *KO* animals over time, and not between *WT* and *Sod1* animals and that likely reflect the consequences of the absence of CSN on the gait: the Max Contact Area, the Print Width Mean, and the Print Area Mean (*WT* vs *KO*, Supplementary Table 1, Slope). We also identified 15 parameters that were significantly altered in *Sod1* mice compared to *WT* mice during disease progression (Supplementary Table 1, Slope). Amongst them, 10 were also significantly different in *Sod1* vs *KO/Sod1* animals (Supplementary Table 1, Slope and Fig. 4). Of those, two parameters showed opposite effects between absence of *Fezf2* and overexpression of *Sod1^G86R^*, such as the Print Position of the Right and Left Paws (Fig. 4k,l; Inter-limb coordination parameter), and eight parameters showed instead a genuine amelioration of the *Sod1* phenotype (*WT* vs *Sod1*) by the absence CNS: the Stand Index (Fig. 4c; temporal parameter), the Max Contact Max Intensity Mean (Fig.4d; spatial parameter) and the Mean Intensity Mean (Fig. 4e; spatial parameter) the Print Length Mean (Fig. 4f; spatial parameter), the Max Intensity Mean (Fig. 4g; spatial parameter) and the Mean Intensity Mean (Fig. 4h; spatial parameter), the Max Intensity of the 15 Most Intense Pixels Mean (Fig. 4i; spatial parameter) and the Number of Steps (Fig. 4j; run characterization parameter). Taken in its whole, this analysis indicates that absence of CSN, *per se*, modifies a small set of spatial gait parameters, but ameliorates a larger set of spatial, temporal, inter-limb coordination and run characterization parameters that are affected in *Sod1^G86R^* animals as disease progresses. Together, the data confirm the beneficial effects of absence of CSN and other SubCerPN on motor impairment.

### Absence of CSN minimizes spinal MN loss and neuromuscular junction denervation without modifying other pathological hallmarks of ALS

To better understand how absence of SubCerPN could be beneficial in the neurodegenerative context of ALS, we further investigated the spinal cord of end-stage animals, for its relevance to ALS but also as the target of CSN, a subpopulation of SubCerPN. To this aim, we ran series of immunolabellings on coronal sections of lumbar spinal cords harvested from end-stage animals and their age-matched control littermates (Fig. 5 and Supplementary Fig. 1). Staining for the astrogliosis and microgliosis markers GFAP and IBA1 revealed reactive gliosis in the lumbar spinal cord of end-stage *Sod1* and *KO/Sod1* animals (Supplementary Fig. 1), but no difference could be observed between these two genotypes. Labelling of the autophagy marker P62/SQTM1 revealed a healthy, homogeneous cytoplasmic labelling in both *WT* and *KO* mice, and large, stellate-like inclusions in the spinal cords of *Sod1* and *KO/Sod1* mice (Supplementary Fig. 1), with similar occurrence and intensity between the two genotypes. Immunolabelling and counting of the ChAT-positive motor neurons (MN) present in the ventral horn of the lumbar spinal cord revealed a mild decrease of the number of MN somas by disease end-stage in *Sod1* mice (Fig. 5a,b), as we previously reported [42,59]. Interestingly, *KO/Sod1* animals maintained a significantly bigger pool of ChAT-positive MN compare to *Sod1* animals (8.456 ± 0.49 vs 6.152 ± 0.57; p = 0.0306; Fig. 5a,b). Spinal MN loss in ALS follows a Wallerian degeneration with an initial denervation of the neuromuscular junctions (NMJ), a consecutive retrograde deterioration of the axons and a final shrinkage and loss of the somas [23]. Thus, to further test whether the mitigation of the motor impairments displayed by *KO/Sod1* mice in comparison with the *Sod1* mice reflected the state of the NMJ, we labelled the NMJ of the tibialis anterior muscle of end-stage *Sod1* and *KO/Sod1* mice and their aged-matched *WT* and *KO* littermates (Fig. 5c). Rating of the NMJ integrity into three categories, innervated, partly denervated and fully denervated, followed by statistical analyses indicated that *KO/Sod1* animals had twice more innervated NMJ than *Sod1* animals by disease end-stage (23.8 ± 4.87 % vs 11.34 ± 3.12 %; p = 0.0239; Fig. 5d). As a whole, the data show that absence of cortical afferences partly protects NMJ denervation and Wallerian degeneration of the MN, and suggest that this partial protection may be independent from local gliosis and altered autophagy.

**Fig. 5.**
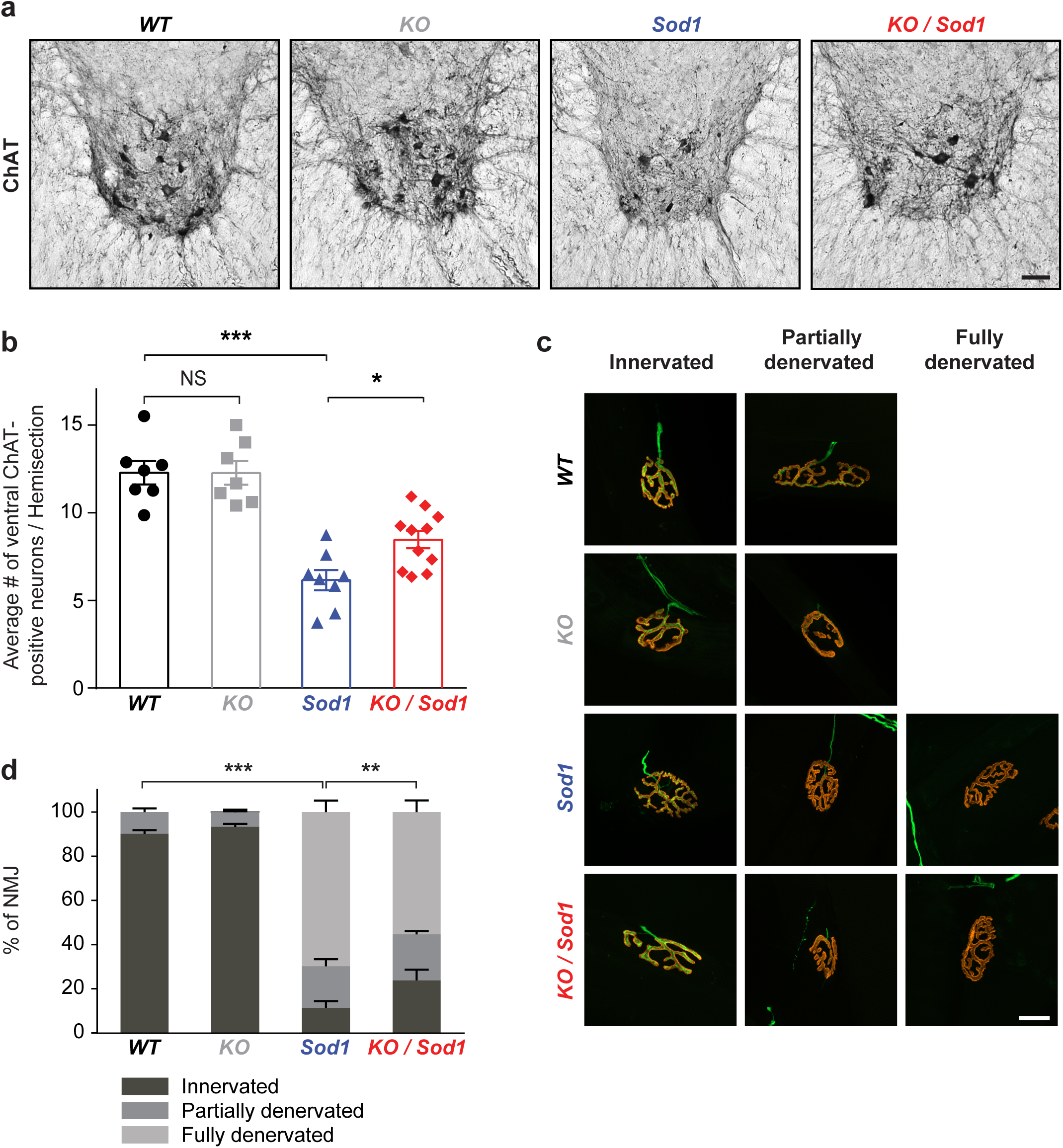
Absence of CSN partially prevents degeneration of the MN cell bodies and NMJ dismantlement. **a** Representative immunohistochemistry images of the ventral horn of the lumbar spinal cord from end-stage *Sod1* and *KO/Sod1* mice and their aged-matched *WT* and *KO* littermates, showing Choline AcetylTransferase-(ChAT-) positive neurons. **b** Bar graph representing the average number of ventral ChAT-positive neurons per lumbar spinal cord hemi-section; 1-way ANOVA; N = 6 *WT*, 6 *KO*, 8 *Sod1*, 10 *KO/Sod1*. **c** Representative maximum intensity projection images of z-stacks of typically innervated, partly or fully denervated neuromuscular junctions (NMJ) from end-stage *Sod1* and *KO/Sod1* mice and their aged-matched *WT* and *KO* littermates. **d** Bar graph representing the average proportions of innervated (dark grey), partly denervated (medium grey) and fully denervated (light grey) NMJ for each genotype; 2-way ANOVA followed by Tukey multiple comparisons test; N = 6 animals per genotype. ** p < 0.05, ** p < 0.01, *** p < 0.001, NS: non-significant; Scale bar: 100 µm in **a,** 20 µm in **c**.

### Absence of CSN positively impacts hyperreflexia but not spasticity

In Humans, degeneration or lesion of the corticospinal tract results in appearance of the upper motor neuron syndrome [52], a series of symptoms that include muscular weakness, decreased motor control, altered muscle tone, hyperreflexia, including spasticity, and clonus [31]. Most ALS patients suffer from spasticity with different degrees of severity [63]. Spasticity is a painful manifestation, defined by J.W. Lance as “a motor disorder characterised by a velocity-dependent increase in tonic stretch reflexes (muscle tone) with exaggerated tendon jerks, resulting from hyperexcitability of the stretch reflex, as one component of the upper motoneuron syndrome” [34]. In ALS patients, hyperreflexia can be measured as the increase of the short latency Hoffman’s reflex, or H-reflex [61]. In rodents, H-reflex *per se*, or the ratio of amplitudes of H-reflex to compound muscle action potentials (H/M ratio) were shown to increase with spasticity [3,47], and long lasting tail muscle activity following cutaneous stimulation was proposed as a simple method to assess spasticity in awake animals [3]. We reasoned that the mouse line that we generated could contribute to assess the role of CSN in the modulation of spinal network excitability involved in hyperreflexia and spasticity, in an ALS-related context.

First, we ran electromyographic analyses on pre-symptomatic mice aged of 80 days (left leg) and 105 days (right leg) in order to detect H-reflex in anaesthetized mice by stimulation of the sciatic nerve and recording in the abductor digiti minimi muscle (Fig. 6a). To verify that the recorded wave that followed the M wave was an H-reflex and not a confounding F wave, known to exhibit similar latencies, we verified that it disappeared with increased stimulus intensity frequency [26] (Supp. Fig. 2). We could detect an H-reflex in fractions of *Sod1* and *KO/Sod1* groups of animals, but never in the *WT* or *KO* animals (Fig. 6a,b). This indicates that, under our experimental conditions, H-reflex could develop only in *Sod1^G86R^* animals, independently of the presence of CSN, and that, counter-intuitively, absence of CSN, *per se*, was not sufficient to trigger an H-reflex (Fig. 6a). Next, we evaluated the proportions of pre-symptomatic animals presenting an H-reflex in either one of the two legs tested (Fig. 6b). No significant difference between *Sod1* and *KO/Sod1* could be seen at this age. However, measurement of the H/M ratio amongst the animals displaying an H reflex showed a significant reduction in *KO/Sod1* animals compared to their *Sod1* littermates (Fig. 6c). This suggests that at pre-symptomatic ages, absence of CSN limits hyperreflexia.

**Fig. 6.**
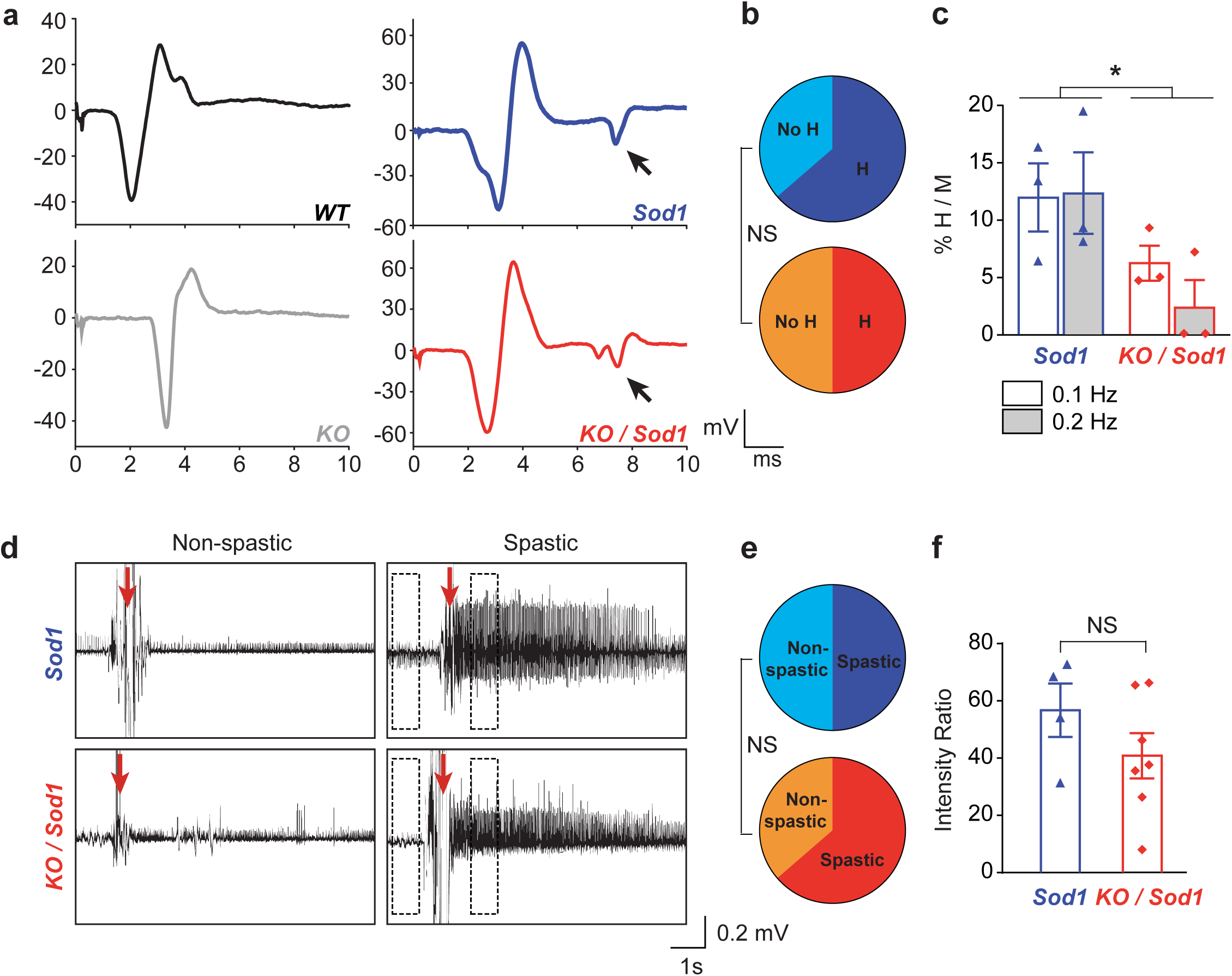
Absence of CSN minimises pre-symptomatic hyperreflexia but end-stage spasticity. **a** Representative EMG traces of the muscular response of the abductor digiti minimi muscle upon sciatic nerve stimulation. Arrows indicate H-reflex in pre-symptomatic *Sod1* and *KO/Sod1* mice; N = 6 *WT*, 6 *KO*, 8 *Sod1*, 10 *KO/Sod1*. **b** Pie charts representing the percentage of animals with an H-reflex amongst *Sod1* (top, blue) and *KO/Sod1* (bottom, red) animals; Fischer exact test. **c** Graph bar representing the averaged ratios of the amplitude of the H and M wave upon repeated stimulations at 0.1 and 0.2 Hz, in *Sod1* and *KO/Sod1* mice; N = 3 *Sod1* and 3 *KO/Sod1*; Two-way ANOVA. **d** Representative EMG recordings of the tail muscle upon stimulation of end-stage, fully paralyzed and awake *Sod1*. Spasticity-related tail long-lasting reflex (LLR, right panels) can be observe upon stimulation (red arrow) in subgroups of animals. **e** Pie charts representing the percentage with a LLR amongst *Sod1* (top, blue) and *KO/Sod1* (bottom, red) animals; Fischer exact test; N = 8 *Sod1*, 11 *KO/Sod1*. **f** Signal-to-noise intensity ratios of the LLR calculated from measurements made in indicates doted boxes in **d**; Student unpaired t-test; N = 4 *Sod1* and 7 *KO/Sod1*; * p < 0.05.

A more integrated assessment of spasticity is the electromyographic evaluation of the tail long lasting reflex (LLR) in awake and fully paralyzed animals prior to harvesting (Fig. 6d-f) [3,15,20]. At disease end stage, the *Sod1* and *KO/Sod1* groups of animals could be subdivided into spastic and non-spastic, but the relative proportions of each was not significantly different between the two genotypes (Fig. 6e). Among spastic animals, intensity of the response to the stimulation was not significantly different between the *Sod1* and *KO/Sod1* groups (Fig. 6f). The data suggest that absence of CSN is not sufficient to prevent the manifestation of spasticity in end-stage animals.

Spasticity of end-stage *Sod1^G86R^* and *SOD1^G37R^* mouse models of ALS has previously been linked to serotonergic neuron degeneration in the raphe nuclei [15,20]. We thus tested whether absence of CSN and other SubCerPN could affect the population of TPH2-positive neurons present in the raphe nuclei over the course of the disease in *Sod1* and *KO/Sod1*, compared to their *WT* and *KO* littermates (Fig. 7). As previously reported [15], end-stage *Sod1* mice displayed a significant loss of TPH2-positive neurons compared to *WT* (Fig. 7b,c). Similarly, we observed a significant loss of TPH2-positive neurons in end-stage *KO/Sod1* mice compared to *WT*, but no significant difference between *Sod1* and *KO/Sod1* animals (Fig. 7b,c). At younger ages, no significant difference could be detected across the different groups of mice. Thus, the data suggest that the parallel loss of serotonergic neurons in end-stage *Sod1* and *KO/Sod1* mice could, at least in part, contribute to the emergence of LLR in end-stage animals, and that absence of CSN and other SubCerPN does not affect serotonergic neurons survival.

**Fig. 7.**
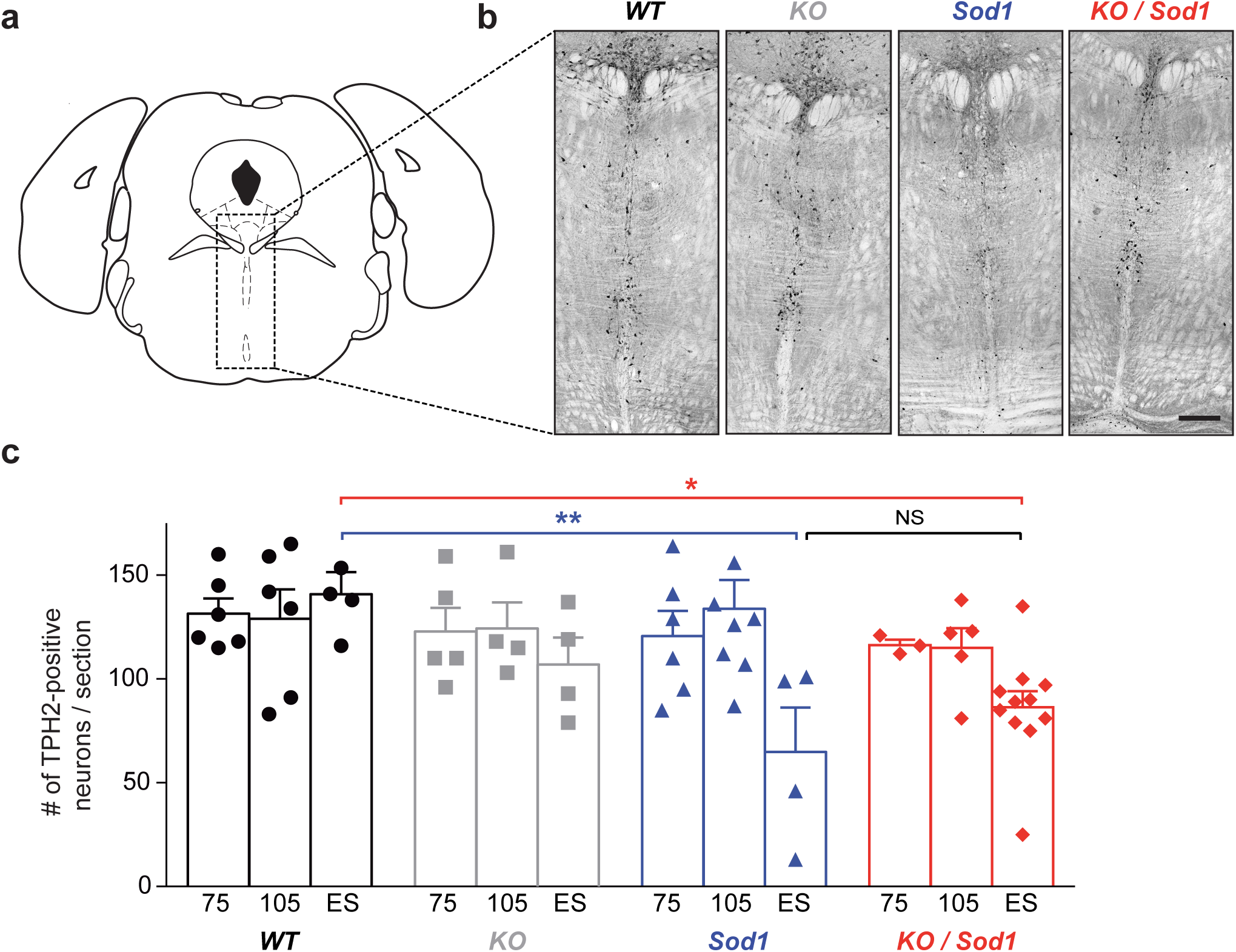
Absence of SubCerPN does not prevent the late loss of serotonergic neurons in the raphe nuclei. **a** Schematic of the coronal section selected for TPH2-positive neurons labelling and counting. **b** Representative images of TPH2 immunoreactivity in the brainstem (dorsal and median raphe) of end stage mice. **c** Bar graph representing the average number of TPH2-positive neurons, over time (75 days, 105 days and ES), in *WT* (N = 6, 6 and 4 respectively), *KO* (N = 5, 4 and 4 respectively), *Sod1* (N = 6, 8 and 4 respectively) and *KO/Sod1* (N = 3, 5 and 11 respectively) mice; 2-way ANOVA followed by Tukey multiple comparisons test; * p < 0.05 and ** p < 0.01; Scale bar 300 µm.

Altogether, the data indicate that hyperreflexia evidenced by an increase of the H/M ratio arises pre-symptomatically in *Sod1^G86R^* mice, and that absence of CSN decreases this particular feature of the upper motor neuron syndrome. Contrastingly, spasticity, evidenced by emergence of the LLR in end-stage *Sod1^G86R^* animals, does not appear to be modulated by the presence or absence of CSN and other SubCerPN, but could, as already demonstrated [15,20], rather arise from the loss of another descending control onto spinal networks, the serotonergic neurons of the raphe nuclei. In mouse models of ALS, H/M ratio may thus represent a better readout of the upper motor neuron syndrome than tail LLR.

## Discussion

In the current study, we sought to test the corticofugal hypothesis of ALS in a well-defined mouse model of the disease. We generated a mouse line overexpressing the *Sod1^G86R^* transgene, a condition sufficient to induce rapidly progressing motor deterioration, premature death and histopathological hallmarks of ALS, but also deficient for the gene *Fezf2* and thus lacking all SubCerPN, the major contributor to the corticofugal projections. Importantly, absence of SubCerPN also implied absence of CSN, a situation that challenged the clinical and histopathological definitions of ALS as the combined degeneration of both MN and CSN. In comparison with single transgenic *Sod1* animals, the resulting double mutant *KO/Sod1* mice presented a delayed disease onset, extended survival, along with improved clinical conditions during disease duration as assessed by decreased weight loss and improved motor performances. In addition, *Sod1* animals presented pre-symptomatic hyperreflexia, a classical feature of the upper motor neuron syndrome, that was also found limited by the absence of SubCerPN and CSN. Together these findings support the corticofugal hypothesis of ALS and may help refocusing pre-clinical research to the cerebral cortex and its neuronal populations.

### Contribution of the cerebral cortex and its outputs to ALS

While the origin of ALS remains a debated question [54], recent evidences from neurophysiological and pathological studies conducted on patients [6,67] are nourishing a revival of interest for Charcot’s initial view of the disease as a primarily cortical impairment [11]. Indeed, whether disease propagation relies on altered neuronal excitability and subsequent excitotoxicity [67], or on prion-like propagation of misfolded proteins [5], both schools of thought converge to a common cortical origin of ALS and propagation along the corticofugal tracts [18,24]. While several clinical features support a cortical origin of the disease [19], and recent longitudinal imaging analyses suggest a propagation of impairments along the corticofugal tracts, [32,66], the hypothesis cannot be tested in patients. If the corticomotoneuronal system has undergone major modifications with evolution [18], it is important to highlight as well the similarities of excitatory cortical neuron subtypes between Human and mouse [30], and to acknowledge that the highest differences arise from the important evolution of the cortico-cortical projection neurons (commissural and associative/intrahemispheric), as opposed to the corticofugal neurons [30,44]. Indeed, corticothalamic sub-populations and subclasses of SubCerPN, *i.e.* cortico-striatal, cortico-tectal, cortico-pontine and cortico-spinal, are well conserved between mouse and Human (reviewed in [44]), suggesting that, while not perfect, rodents may rather be good models to study the contribution of SubCerPN to ALS, and to directly test the corticofugal hypothesis.

Further supporting the appropriateness of rodents in general and mice in particular to cortical impairment in ALS, is their ability to recapitulate CSN [28,42,50,70,71] or SubCerPN degeneration [22,38]. In that respect, we recently completed a comprehensive spatiotemporal analysis of CSN degeneration in the *Sod1^G86R^* mouse model of ALS, and described a loss of CSN that precedes that of MN and even NMJ denervation. In this model, CSN and MN degenerations are also somatotopically related, suggesting that cortical impairment precedes MN impairment [42]. Of further relevance to Human disease, it is noteworthy that CSN loss also starts before weight loss, pointing to the earliness of cortical impairment in these animals [16,42]. This is further revealed in the present study by the pre-symptomatic occurrence of hyperreflexia in sub-groups of *Sod1* and *KO/Sod1* animals, which brings here a functional evidence of early impairment of the CSN. Thus, the rodent models in general and *Sod1^G86R^* mice in particular seem appropriate to assess the contribution of CSN and other SubCerPN to ALS, and to test whether CSN degeneration may be detrimental to downstream MN by lack of protective input, or instead be beneficial by limiting the propagation of a toxic input from the motor cortex.

### Absence of SubCerPN delays onset and extends survival without increasing disease duration

In this study, we sought to assess specifically the contribution of CSN and other SubCerPN to disease onset and progression in mice. To this aim, we crossbred the *Sod1^G86R^* and the *Fezf2^−/−^* mouse lines to generate a model that ubiquitously expresses a mutant of the murine *Sod1* gene [56] but lacks CSN and other SubCerPN [29,48], hence challenging ALS pathological and clinical definitions. Our data indicate that absence of SubCerPN delays disease onset and increases survival without modifying disease duration. The results are highly reminiscent of an elegant study by Thomsen and colleagues who knocked-down mutant *SOD1* transgene in the posterior motor cortex of the *SOD1^G93A^* rat model of ALS [62]. The authors chose an AAV9 virus to selectively transduce neurons. AAV9-SOD1-shRNA injections delayed disease onset and extended survival without affecting disease duration [62]. The similarity of the results between this study and ours is remarkable given the broad differences of the two approaches. First, while in *Sod1^G86R^* mice, CSN degenerate long before MN [42], CSN degeneration is secondary to MN loss in the *SOD1^G93A^* rats [62]. Second, while *Sod1^G86R^* mice were crossed to the *Fezf2 KO* mice to prevent any direct connection between the motor cortex and its targets, the knocked-down of the *SOD1^G93A^* transgene in rats was undertaken to favour the maintenance of a functional pool of CSN. Together, the two sets of data suggest that absence of diseased CSN or instead maintenance of genetically corrected CSN may be equally beneficial, and that *Sod1/SOD1* mutant transgene-expressing CSN may be detrimental to their downstream targets. Yet, it is worth mentioning that AAV9 likely targeted all types cortical excitatory projection neurons and inhibitory interneurons, and could have contributed to correct more broadly the cortical circuit dysfunctions which have been reported in the *Sod1^G93A^* and the *TDP-43^A315T^* mouse models of ALS [33,73], and are reminiscent of the cortical hyperexcitability that characterizes ALS patients [67]. Future cell-type specific genetic ablation experiments could further inform on the contribution of individual populations of neurons and glia to cortical circuit dysfunction and impairment on the downstream targets of the cerebral cortex. Such approaches have allowed to better define the contributions of MN [4], microglia [4,68], astrocytes [69] and muscles [45] to ALS onset and progression. Together, these studies and ours indicate that cerebral cortex and SubCerPN may rather be involved in disease onset, while MN and glia may rather modulate disease progression.

### Lack of SubCerPN attenuates weight loss

Far above the cortico-motoneuronal axis, ALS affects the whole-body physiology and in particular energy metabolism. Weight loss of ALS patients is now acknowledged to start long before motor symptoms appearance, and massive weight loss negatively correlates with survival [16,17,51]. Energy homeostasis dysfunction is likely to arise, at least partly, from impairment of the hypothalamus, a subcortical structure implicated in food intake and weight control [64]. Clear hypothalamic atrophy is present in ALS patients, as well as pre-symptomatic gene carriers, and hypothalamic volume has been negatively correlated to disease onset [25]. While the hypothalamus is known to receive indirect control from the prefrontal cortex via the amygdala, it may also, at least in the rat, be under direct control of the medial prefrontal cortex via its layer V SubCerPN [55]. It is thus possible that the cerebral cortex and its corticofugal outputs also contribute, at least partly, to weight loss in ALS. Such a scenario would require further investigation of the hypothalamus and the energy metabolism of mice that lack SubCerPN.

### Absence of CSN limits hyperreflexia, MN loss and NMJ dismantlement

Absence of SubCerPN in *Sod1^G86R^* mice is beneficial to MN connection to their muscular targets, as assessed by the improvement of the motor performances, during the course of the disease, of double mutant mice compared to single mutant mice, and by increased numbers of MN and innervated NMJ at disease end-stage in *KO/Sod1* versus *Sod1* animals. Our data indicate that this mitigation of signs of MN degeneration was accompanied by amelioration of hyperreflexia, a feature of the upper motor neuron syndrome [31,61]. We evaluated hyperreflexia by detecting the H-reflex in the abductor digiti minimi muscle upon stimulation of the sciatic nerve, and quantifying the H/M ratio, as previously reported [3,47]. Absence of H-reflex recording in *WT* animals does not rule out the existence of a small amplitude H-reflex in these animals, below the detection threshold of our recording set-up, as already observed [47]. In contrast, presence of an H-reflex in a subgroup of pre-symptomatic *Sod1* animals suggest that hyperreflexia may take place in this mouse model of ALS, long before appearance of the motor symptoms. Importantly, this correlates with the early pre-symptomatic degeneration of the CSN that we reported in this mouse line [42]. Absence of H-reflex from *KO* animals suggests that developmental absence of CSN and other SubCerPN may have been compensated by other supraspinal controls or by spinal network rearrangements, or both, preventing the emergence of hyperreflexia. Finally, decreased H/M ratios in *KO/Sod1* compared to *Sod1* animals indicate a decreased hyperreflexia, and a possible mitigation of the upper motor neuron syndrome in absence of CSN and other SubCerPN. Contrastingly, spasticity, assessed by emergence of the long lasting reflex (LLR) of the tail muscle of fully paralysed end-stage mice is neither prevented nor even slightly modulated by absence of SubCerPN and CSN. However, evaluation of the LLR can only be performed in paralyzed animals, and our data do not rule out that spasticity, while present in *KO/Sod1* mice, might have arisen later. Recent studies support the role of serotonergic neurons in the emergence of spasticity in ALS [15,20,37,49]. Our data confirm the loss of TPH2-positive serotonergic neurons in the raphe nuclei of end-stage *Sod1^G86R^* mice, independently from the presence or absence of CSN. However, loss of serotonergic neurons is a late event in *Sod1^G86R^* mice, and for technical reasons tail LLR could not be tested earlier (*i.e.* before full paralysis of the animals). It is thus difficult to estimate when LLR starts, and whether it could potentially, at earlier time points, reflect CSN degeneration. Importantly, because no loss of serotonergic neurons could be detected when we recorded H-reflex, it is unlikely that in the *Sod1^G86R^* mouse model of ALS emergence of hyperreflexia, as assessed by H-reflex occurrence, could arise from serotonergic neuron degeneration. In agreement with previous reports [15,20], our data suggest that tail LLR may rather reflect serotonergic neuron loss than CSN loss. We thus propose to use H-reflex and H/M ratios as readouts of CSN degeneration and upper motor syndrome in ALS mouse models.

### Perspectives

In this study, we demonstrated that absence of SubCerPN was beneficial in a mouse model of ALS, suggesting that major corticofugal projections may be detrimental to their downstream targets in a context of ALS. Yet, because the strategy we employed prevented the development of a broad neuronal population, it is possible that compensations by other nuclei have occurred. For instance, genetic ablation of the corticospinal tract in the *Celsr3/Emx1* mice was shown to induce both increased numbers of rubrospinal projections and of terminal ramification of monoaminergic axons [27]. Genetic ablation of adult CSN and SubCerPN would thus be particularly informative to better understand how these neurons may contribute to disease onset and progression. While our study clearly demonstrated a detrimental role of CSN and other SubCerPN in a mouse model of ALS, it did not inform on the nature of this toxic effect, *i.e.* whether it could arise from altered excitability and downstream glutamatergic excitotoxicity, or from potential propagation of misfolded SOD1 protein, or both. Selective silencing of, or selective transgene excision from the SubCerPN, along with a deep mechanistic analysis of the transcriptomic modifications that accompany CSN dysfunction and degeneration in ALS [42], may in the future provide a better understanding of the role of the cerebral cortex and its outputs to ALS, and potentially unravel new therapeutic targets.

## Acknowledgments

The work has been supported by a European Research Council (ERC) starting grant #639737, a Marie Curie career integration grant #618764, an “Association Française contre les Myopathies” (AFM)-Telethon trampoline grant #16923 and a Neurex grant to CR, PhD fellowships from the French Ministry of Research to TB and CB, followed by the “Fondation pour la Recherche sur le Cerveau” (FRC) to TB and the “Association de Recherche sur la Sclérose Latérale Amyotrophique” (ARSLA) to CB. JCZ was supported by a post-doctoral fellowship from the AFM-Telethon. The authors are extremely thankful to Véronique Marchand-Pauvert, Pascal Branchereau, Pierre Veinante and Luc Dupuis for critical reading of the manuscript, and insightful comments.

---

**Supplementary Fig. 1.**
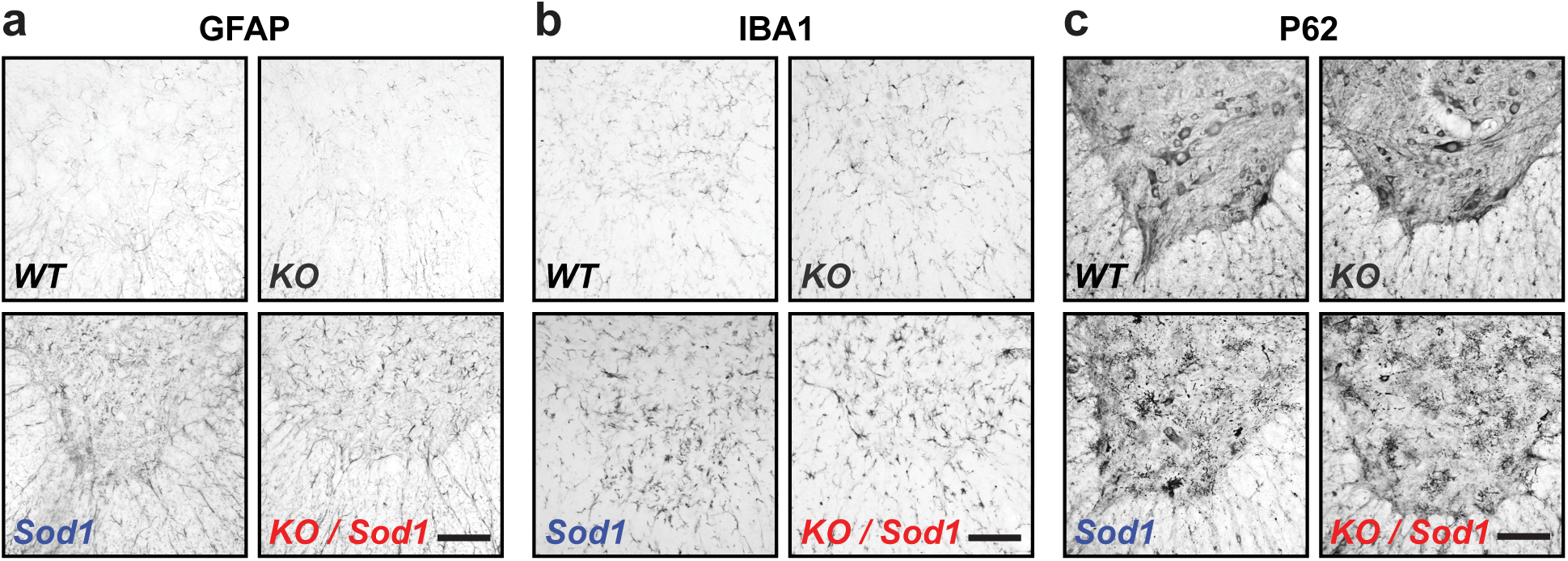
Absence of SubCerPN does not modify the hallmarks of ALS pathology in the spinal cord. **a-c** Representative immunostaining images of the ventral horn of the lumbar spinal cord from end-stage *Sod1* and *KO/Sod1* mice and their age-matched *WT* and *KO* littermates. *Sod1* and *KO/Sod1* mice displayed increased GFAP (**a**) and IBA1 (**b**) immunoreactivity, as well as P62-positive aggregates (**c**) compare to *WT* and *KO* mice. N = 6 animals per genotype; Scale bar 100 µm.

**Supplementary Fig. 2.**
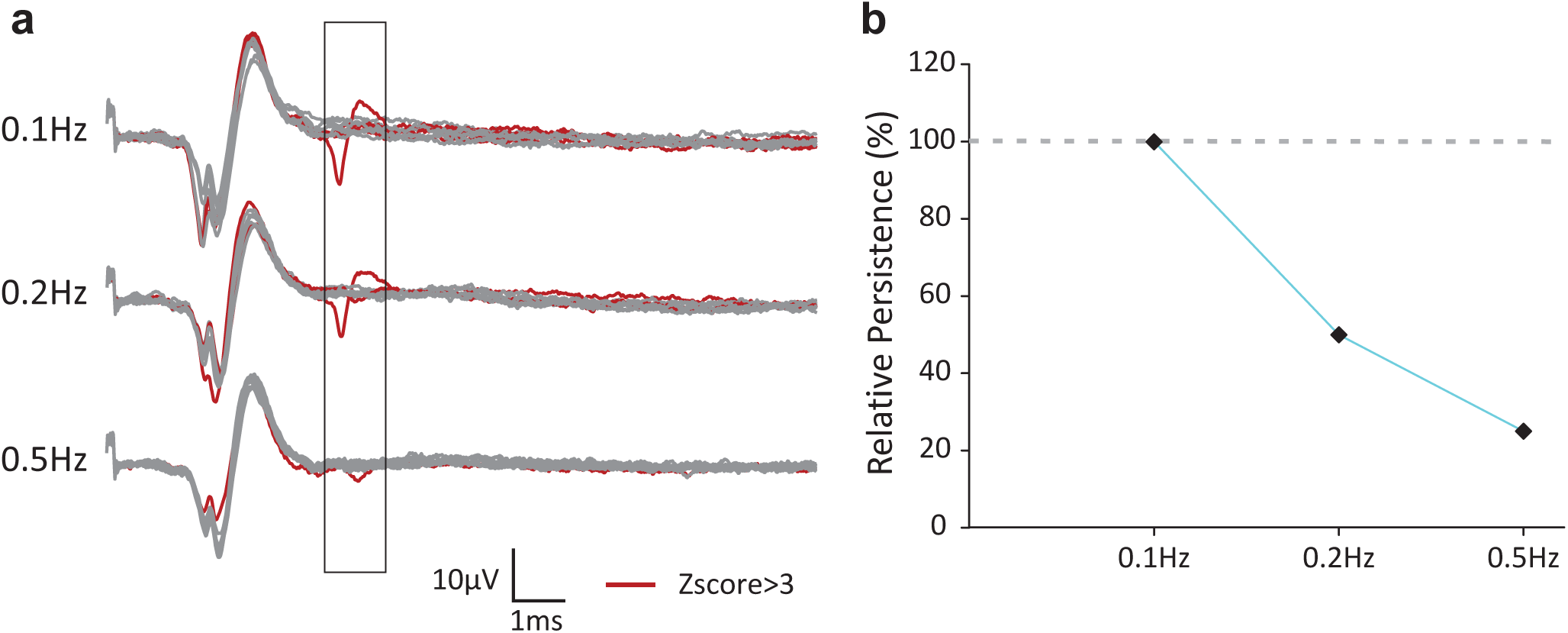
Decreased persistence of H-Reflex with increased stimulation frequency. **a** Representative EMG traces of the muscular response in the abductor digiti minimi muscle upon sciatic nerve stimulation resulting from 10 stimulations at 0.1, 0.2 and 0.5Hz. The traces in red represent signals containing a significant response (Z-score ≥ 3) in the measured interval (squared in black). **b** Representative averaged values of the persistence of the H-Reflex at different stimulation frequencies in relation to the 0.1Hz stimulation.

**Supplementary Table 1.** Linear regression analysis of the different motor tests and CatWalk parameters. List of parameters analysed by linear regression with either *i)* no covariate (left columns), *ii)* weight only (middle columns), *iii)* weight and speed as covariates (right columns). “4 Paws” parameters represent averages of the data of each individual paw. Analyses were performed when experiments were initiated (Origin), as well as during the whole course of the experiment (Slope); * p < 0.05; ** p < 0.01; *** p < 0.001; NS = non-significant.

