## Supplementary Table for "Absence of subcerebral projection neurons delays disease onset and extends survival in a mouse model of ALS"

**Burg et al., Supplementary Table 1 :** Linear regression analyses of the different motor tests and CatWalk parameters

| Parameters | No Covariate | | | | | | Covariate = Weight | | | | | | Covariates = Weight + Speed | | | | | |
|---|---|---|---|---|---|---|---|---|---|---|---|---|---|---|---|---|---|---|
| | ORIGIN | | | SLOPE | | | ORIGIN | | | SLOPE | | | ORIGIN | | | SLOPE | | |
| | WT vs Sod1 | WT vs KO | Sod1 vs KO / Sod1 | WT vs Sod1 | WT vs KO | Sod1 vs KO / Sod1 | WT vs Sod1 | WT vs KO | Sod1 vs KO / Sod1 | WT vs Sod1 | WT vs KO | Sod1 vs KO / Sod1 | WT vs Sod1 | WT vs KO | Sod1 vs KO / Sod1 | WT vs Sod1 | WT vs KO | Sod1 vs KO / Sod1 |
| Grip | NS | *** | * | *** | NS | *** | *** | ** | * | *** | NS | *** | | | | | | |
| Rotarod | ** | NS | NS | *** | NS | *** | *** | NS | *** | *** | NS | ** | | | | | | |
| Inverted Grid | NS | *** | *** | *** | NS | *** | *** | *** | NS | *** | NS | *** | | | | | | |
| Run Average Speed | *** | ** | * | NS | NS | * | | | | | | | | | | | | |
| 4 Paws Stand Mean | *** | *** | * | NS | NS | NS | NS | NS | NS | NS | NS | NS | NS | NS | NS | NS | NS | NS |
| 4 Paws Stand Index Mean | * | NS | * | *** | NS | ** | NS | NS | NS | NS | NS | NS | NS | NS | NS | *** | * | ** |
| 4 Paws Max Contact At Mean | NS | NS | NS | NS | * | NS | NS | NS | NS | NS | NS | NS | NS | NS | NS | NS | NS | NS |
| 4 Paws Max Contact Area | NS | * | NS | NS | * | NS | NS | NS | NS | * | ** | * | NS | NS | NS | NS | *** | * |
| 4 Paws Max Contact Max Intensity Mean | NS | * | NS | * | * | * | NS | NS | NS | *** | ** | ** | NS | NS | NS | *** | ** | ** |
| 4 Paws Max Contact Mean Intensity Mean | NS | * | NS | NS | *** | NS | NS | NS | NS | ** | ** | * | NS | NS | NS | ** | *** | * |
| 4 Paws Print Length Mean | NS | NS | NS | ** | ** | ** | NS | NS | NS | *** | NS | ** | NS | NS | NS | *** | NS | ** |
| 4 Paws Print Width Mean | NS | NS | NS | NS | NS | NS | NS | NS | NS | NS | * | * | NS | NS | NS | NS | ** | NS |
| 4 Paws Print Area Mean | NS | * | NS | NS | * | NS | NS | NS | NS | NS | ** | * | NS | NS | NS | NS | *** | * |
| 4 Paws Max Intensity At Mean | NS | * | NS | NS | NS | NS | NS | NS | NS | NS | NS | NS | NS | NS | NS | NS | NS | NS |
| 4 Paws Max Intensity Mean | NS | * | NS | * | * | * | NS | NS | NS | ** | ** | ** | NS | NS | NS | ** | ** | ** |
| 4 Paws Min Intensity Mean | NS | NS | * | NS | NS | NS | NS | NS | NS | NS | NS | NS | NS | NS | NS | NS | NS | NS |
| 4 Paws Mean Intensity Mean | NS | * | * | NS | *** | NS | NS | NS | NS | ** | *** | * | NS | NS | NS | ** | *** | * |
| 4 Paws Mean Intensity of the 15 most intense pixels Mean | NS | ** | * | * | ** | * | NS | NS | NS | ** | *** | ** | NS | NS | NS | ** | *** | ** |
| 4 Paws Swing Mean | ** | * | NS | NS | NS | NS | NS | NS | NS | NS | NS | NS | NS | NS | NS | NS | NS | NS |
| 4 Paws Swing Speed Mean | ** | *** | *** | NS | NS | NS | ** | NS | NS | NS | NS | NS | NS | NS | NS | NS | NS | NS |
| 4 Paws Stride Length Mean | NS | * | * | NS | NS | * | ** | NS | ** | NS | NS | NS | NS | NS | NS | NS | NS | NS |
| 4 Paws Step Cycle Mean | *** | *** | * | NS | NS | NS | NS | NS | NS | NS | NS | NS | NS | NS | NS | NS | NS | NS |
| 4 Paws Duty Cycle Mean | NS | NS | NS | * | NS | * | NS | NS | * | NS | NS | NS | NS | NS | NS | NS | NS | NS |
| 4 Paws Single Stance Mean | *** | ** | ** | NS | NS | NS | NS | NS | NS | NS | NS | NS | * | NS | NS | NS | NS | NS |
| 4 Paws Initial Dual Stance Mean | *** | *** | NS | NS | NS | NS | * | NS | NS | NS | NS | NS | NS | NS | NS | NS | NS | NS |
| 4 Paws Terminal Dual Stance Mean | *** | *** | * | NS | NS | NS | * | NS | NS | NS | NS | NS | NS | NS | NS | NS | NS | NS |
| 4 Paws Body Speed Mean | * | ** | ** | NS | NS | * | ** | NS | * | NS | NS | NS | NS | NS | NS | NS | NS | NS |
| 4 Paws Body Speed Variation Mean | NS | NS | *** | NS | NS | NS | NS | NS | ** | NS | NS | NS | NS | NS | NS | NS | NS | NS |
| Step sequence number of Patterns | NS | ** | * | NS | NS | NS | *** | NS | *** | NS | NS | NS | NS | NS | *** | NS | NS | NS |
| Step sequence CA | NS | NS | NS | NS | NS | NS | NS | NS | NS | NS | NS | NS | NS | NS | NS | NS | NS | NS |
| Step sequence CB | NS | NS | NS | NS | NS | NS | NS | NS | NS | NS | NS | NS | NS | NS | NS | NS | NS | NS |
| Step sequence AA | *** | NS | *** | NS | NS | NS | NS | NS | NS | NS | NS | NS | NS | NS | ** | NS | NS | NS |
| Step sequence AB | *** | * | NS | NS | NS | NS | NS | NS | * | NS | NS | NS | ** | NS | *** | NS | NS | NS |
| Step sequence RA | NS | NS | NS | NS | NS | NS | * | NS | NS | ** | NS | NS | ** | NS | NS | ** | NS | NS |
| Step sequence RB | NS | * | NS | NS | NS | NS | NS | NS | NS | NS | NS | NS | NS | NS | NS | NS | NS | NS |
| Step sequence Regularity index | NS | NS | NS | NS | NS | NS | * | NS | NS | * | NS | NS | * | NS | ** | * | NS | NS |
| BOS Frontpaws | NS | NS | *** | NS | NS | NS | NS | NS | NS | NS | NS | NS | NS | NS | NS | NS | NS | NS |
| BOS Hindpaws | NS | NS | *** | NS | NS | * | NS | NS | NS | *** | NS | NS | NS | NS | NS | ** | NS | NS |
| Duration Mean | *** | *** | * | NS | NS | NS | *** | NS | NS | NS | NS | NS | NS | NS | NS | * | NS | NS |
| Average Speed Mean | * | ** | *** | NS | NS | * | ** | NS | NS | NS | NS | NS | NS | NS | NS | NS | NS | NS |
| Maximum variation Mean | NS | NS | NS | NS | NS | NS | NS | NS | NS | NS | NS | NS | NS | NS | NS | NS | NS | NS |
| Number of Steps | NS | * | * | NS | NS | NS | *** | NS | *** | NS | NS | NS | *** | NS | *** | ** | NS | ** |
| Print position Right paws Mean | NS | *** | *** | * | * | NS | *** | NS | *** | *** | NS | *** | *** | NS | *** | *** | NS | *** |
| Print position Left paws Mean | NS | *** | *** | *** | NS | NS | *** | *** | *** | *** | NS | *** | *** | *** | *** | *** | * | *** |
| Support Zero | NS | NS | NS | NS | NS | NS | NS | NS | NS | NS | NS | NS | NS | NS | NS | NS | NS | NS |
| Support Single | NS | NS | NS | ** | NS | ** | NS | NS | NS | NS | * | NS | ** | NS | NS | *** | NS | NS |
| Support Diagonal | NS | NS | NS | NS | NS | NS | ** | NS | NS | NS | NS | NS | NS | NS | NS | NS | NS | NS |
| Support Girdle | NS | NS | NS | NS | NS | NS | NS | NS | NS | NS | NS | NS | NS | NS | NS | NS | NS | NS |
| Support Lateral | NS | *** | NS | NS | NS | NS | NS | NS | *** | NS | NS | NS | NS | NS | *** | NS | NS | NS |
| Support Three | NS | NS | NS | NS | NS | * | NS | NS | * | NS | NS | NS | NS | NS | NS | NS | NS | NS |
| Support Four | * | *** | NS | NS | NS | NS | * | NS | * | NS | NS | NS | NS | NS | NS | NS | NS | NS |
